# Porcine circovirus 2 uses a multitude of weak binding sites to interact with heparan sulfate, and the interactions do not follow the symmetry of the capsid

**DOI:** 10.1101/370411

**Authors:** Sonali Dhindwal, Bryant Avila, Shanshan Feng, Reza Khayat

## Abstract

Porcine circovirus 2 is the smallest pathogenic virus capable of autonomous replication within its host. Infections result in immunosuppression and subsequent death of the host, and are initiated via the attachment of the PCV2 icosahedral capsid to heparan sulfate and chondroitin sulfate B glycosaminoglycans on the cell surface. However, the underlying mechanism of structural recognition remains to be explored. Using heparin, a routinely used analog of heparan sulfate, we demonstrate that increasing lengths of heparin exhibit greater affinity towards PCV2. Our competition assays indicate that dextra sulfate (8kDa) has higher affinity than heparin (12kDa), chondroitin sulfate B (41kDa) hyaluronic acid (1.6MDa), and dextran (6kDa) for PCV2. This suggests that polymers high in sulfate content are capable of competing with the PCV2-heparan sulfate interaction, and thus have the potential to inhibit PCV2 infection. Finally, we visualize the interaction between heparin and the PCV2 capsid using cryo-electron microscopy single particle analysis, symmetry expansion, and focused classification. The image reconstructions provide the first example of an asymmetric distribution of heparin on the surface of an icosahedral virus capsid. We demonstrate that each of the 60 capsid subunits that generate the *T*=1 capsid can bind heparin via one of five binding sites. However, not all of the binding sites are occupied by heparin and only one-to two-thirds of the binding sites are occupied. The binding sites are defined by arginine, lysine, and polar amino acids. Mutating the arginine, lysine, and polar amino acids to alanine diminishes the binding capacity of PCV2 to heparin.

**Importance:** It has been demonstrated that porcine circovirus 2 (**PCV2**) attaches to cells via heparan sulfate (**HS**) and chondroitin sulfate B (**CSB**) glycosaminoglycans; however, the underlying structural mechanism describing the HS/CSB recognition by PCV2 remains to be explored. We use cryo-electron microscopy with single particle analysis, symmetry expansion, and focused classification to visualize the interaction between the PCV2 capsid and heparin, an analog of heparan sulfate, to better than 3.6Å resolution. We observe that the interaction between the PCV2 and heparin does not adhere to the icosahedral symmetry of the capsid. To the best of our knowledge, this is the first example where the interaction between heparin and an icosahedral capsid does not follow the symmetry elements of the capsid. Our findings also suggest that anionic polymers such as dextran sulfate may act to inhibit PCV2 infection.

## Introduction

Porcine circovirus (**PCV**) belongs to the *Circovirus* genus of the *Circoviridae* family (1, 2). Members of this family are non-enveloped viruses and possess a circular single stranded DNA (**ssDNA**) genome. Circoviruses are widely distributed in nature and infect terrestrial, avian, and aquatic members of the animal kingdom (3, 4). Three genotypes of PCV have been identified –PCV1, PCV2 and PCV3. PCV1 (1,759nt) was first detected in porcine kidney (PK-15) cell lines, and later found to be a nonpathogenic virus (5, 6). PCV2 (1,767nt to 1,768nt) is morphologically similar to but genetically and antigenically distinct from PCV1, and was isolated from pigs with Postweaning Multisystemic Wasting Syndrome (**PMWS**) (3, 7–10). PMWS, later named to porcine circovirus-associated disease (PCVAD) or porcine circovirus disease (PCVD), culminates in the immunosuppression of the host and death from secondary infection (11–14). Autopsy of infected pigs identifies PCV2 in nearly every tissue, indicating that it has a broad tissue tropism (1, 2, 11, 15). The promiscuous nature of PCV2 is further exhibited by its ability to infect and induce its pathogenic phenotype in rodents and bovine living in the vicinity of infected farms, and BALB/c mice and human cells in the laboratory (3, 4, 16–19). PCV3 (2,000nt) was recently identified and shown to be associated with porcine dermatitis, reproductive failure, and nephropathy syndrome (5, 6, 20).

PCV2 is the smallest pathogenic virus capable of replicating in cells without the need for additional viruses (3, 7–10, 14). Its ^∼^1.7knt ambisense genome encodes for a replicase (ORF1) responsible for the rolling circle replication of the genome, a capsid protein (ORF2) responsible for forming the capsid and enclosing the genome, and an ORF3 and ORF4 which may be responsible for causing cellular apoptosis and the pathogenic nature of PCV2 (5, 11–14, 21–24). PCV2 has been shown to initiate cellular infection via attachment to the glycosaminoglycans (**GAG**s) heparan sulfate and chondroitin sulfate B (25). Heparan sulfate (**HS**) and chondroitin sulfate B (**CSB**) are ubiquitously expressed on mammalian cells and act as attachment factors for a variety of macromolecules such as proteases, chemokines, receptors, and pathogens (26, 27). Heparan sulfate is a 30 to 70 kDa linear polysaccharide (40-300 sugar residues and approximately 20-150nm long) composed of alternating sulfated domains (**NS**) and unsulfated domains (**NA**) (26, 28). The NS domains are composed of three to eight repeating disaccharides of L-Iduronic acid (**IdoA**) and D-Glucosamine (**GlcN**) (**Figure S1**). An NS disaccharide can possess two to three sulfates. The NA domains are composed of two to twelve repeating disaccharides of N-Acetyl-D-Glusomaine (**GlcNAc**) and D-Glucoronic acid (**GlcA**) (**Figure S1**) (29). Thus a single chain of HS is composed of multiple NS and NA domains. The NS domain is similar to the heparin structure, and heparin is routinely used as a reagent to study the interaction between the NS domains and their interacting partners (28). Heparin is not a cellular attachment factor, but stored in the granules of mast cells and released into the vascular system as an anti-inflammatory agent (30). CSB, also a linear polysaccharide, is composed of the repeating disaccharide N-Acetyl-Galactosamine (**GalNAc**) and IdoA. A CSB disaccharide has one to two sulfates (27).

PCV2 has been shown to internalize via two distinct pathways: clathrin-mediated endocytosis in monocytic (31) and dendritic (1) cells, and caveolae-, clathrin and dynamin-independent small GTPase regulated pathways in epithelial cell lines of porcine kidney (PK15), swine kidney (SK) and swine testicles (ST) (3). The PCV2 nucleocapsid has been shown to gain cellular entry by escaping the acidified endosome-lysosome of 3D4/31 cells (31). A bipartite nuclear localization signal (**NLS**) at the N-terminus of the capsid protein has been implicated to guide the nucleocapsid into the nucleus for genome replication (32). The PCV2 replicase (ORF1) recruits the cellular machinery to initiate rolling circle replication of the PCV2 genome in the nucleus (33). The newly synthesized PCV2 capsid protein is transported into the nucleus, via its NLS, for genome encapsidation and assembly of infectious virion. Virion then egress from the cell to initiate another cycle of infection (3, 8, 10).

The PCV capsids are 20nm in diameter, possess icosahedral symmetry, and are composed of 60-copies (*T=1*) of a capsid protein (34, 35). From here forth we use the terminology “capsid subunit” to refer to the capsid protein in the context of the assembled capsid. The crystal structure of the PCV2 virus like particle (**VLP**) visualized the capsid subunit fold to be that of the canonical viral jelly-roll, first observed for the capsid subunit of tobacco bushy stunt virus (35, 36). The subunit can be described as a β-sandwich, or β-barrel, with a right hand twist (**Figure 1A**). The β-sandwich is composed of two β-sheets. One β-sheet is made of the four antiparallel β-strands BIDG, and the second β-sheet is composed of the four antiparallel β-strands CHEF (**Figure 1A**). The interface between the two β-sheets is the hydrophobic core of the protein. Loops connecting the strands generate the surface topography of the VLP. Strands BC, DE, FG and HI are short and surround the 5-fold vertex of the VLP (**Figure 1B**). Loops connecting strands CD, EF and GH are longer. Loop CD from neighboring subunits contact one another across the icosahedral 2-fold axes of symmetry, and loop GH from neighboring subunits contact one another across the icosahedral 3-fold axes of symmetry (**Figure 1B**).

**Figure 1.**
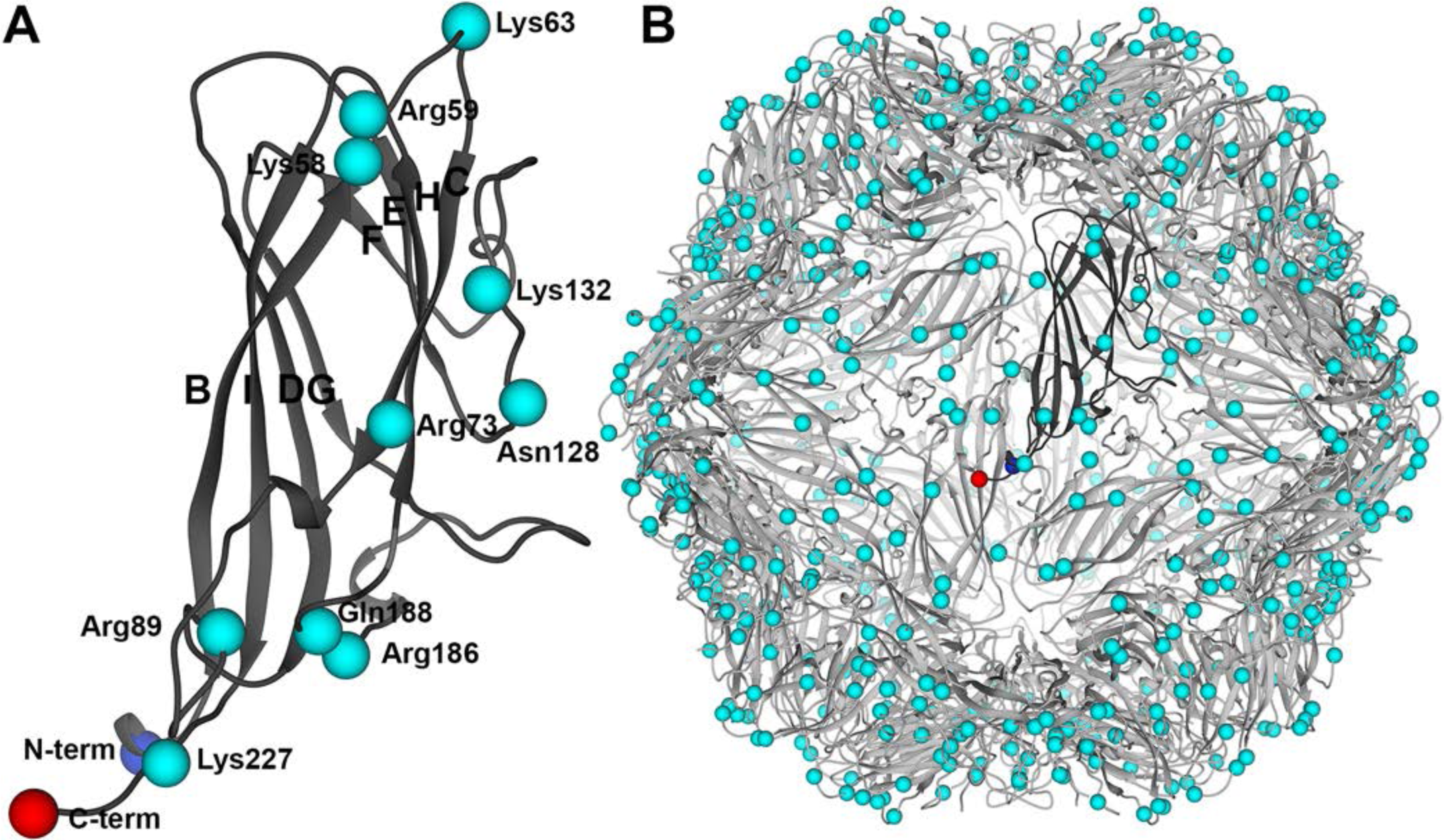
Cartoon depiction of the PCV2 subunit and VLP atomic coordinates. A) Ribbon diagram of the capsid subunit. The β-strands (BCDEFGHI) are labeled according to the sequence in which they occur. The N- and C-termini are shown as a blue and red sphere, respectively. The Cα atoms for basic and polar residues identified as heparin binding residues in this study are depicted as cyan spheres. B) Ribbon diagram of the VLP showing the molecular organization of the subunits. A single subunit is shown in a dark grey color. Images produced with UCSF Chimera (64).

Because the structural mechanism of cellular recognition is not understood, we use combination of cryo-electron microcopy, biochemistry, and site directed mutagenesis to describe this interaction. We use heparin as an analog for studying the interaction between heparan sulfate and PCV2. We ask, does the length of heparin affect its affinity for PCV2, what chemical moieties of GAGs interact with PCV2, where are the heparin binding sites on PCV2, does binding of heparin follow the icosahedral symmetry of the capsid, and does PCV2 undergo a conformational change upon binding of heparin? Our biochemical results demonstrate that longer oligosaccharides of heparin bind with greater affinity to the capsid, and that that the interaction is predominantly driven by the sulfates of the polysaccharide. Our structural results visualize heparin to bind one of five binding sites per capsid subunit, and the interaction to not adhere to the capsid’s icosahedral symmetry. Finally, binding of heparin does not induce the Cα backbone to undergo a conformational change. To our knowledge, this is the first example of a non-symmetric distribution of heparin on an icosahedral virus. Furthermore, our studies provide the first structural study of a member of the *Circovirus* genus interacting with the cellular attachment factor of a cell to initiate infection. The knowledge gained in this study can pave the path for developing molecules to interfere with this interaction and inhibit PCV2 infection.

## Results

### The interaction between PCV2 and heparin is reversible, dependent on the size of heparin, and primarily dictated by sulfates

To study the interaction between heparan sulfate and PCV2 we used an *in vitro* binding assay that involves interacting PCV2 virus like particles (**VLP**) with chromatography sorbent conjugated to 15kDa porcine intestinal mucosa heparin. Heparin is routinely used as an analog of heparan sulfate (**HS**) for studying the interaction between macromolecules and HS (26, 29). This is because the structure of heparin is similar to the NS domain of HS (**Figure S1**). We first determined the concentration of Baculovirus expressed PCV2 VLP (GenBank: ACA51584.1) and time necessary to interact with the sorbent to achieve a robust readout (**Figure 2A**). Two concentrations of VLP were used (370 nM and 92 nM). Maximum binding occurred within 30 minutes for the lower concentration of VLP, and within 3 hours for the higher concentration of VLP. We chose to use the higher concentration of VLP (370 nM) for our subsequent studies because of its improved signal in our binding assays (**Figure S2A).** We then examined if the interaction was reversible by incubating the PCV2 bound sorbent with increasing concentrations of 12kDa porcine intestinal mucosa heparin (**Figure 2B, S2B**). Elution of PCV2 from the sorbent demonstrates that the interaction is reversible.

**Figure 2.**
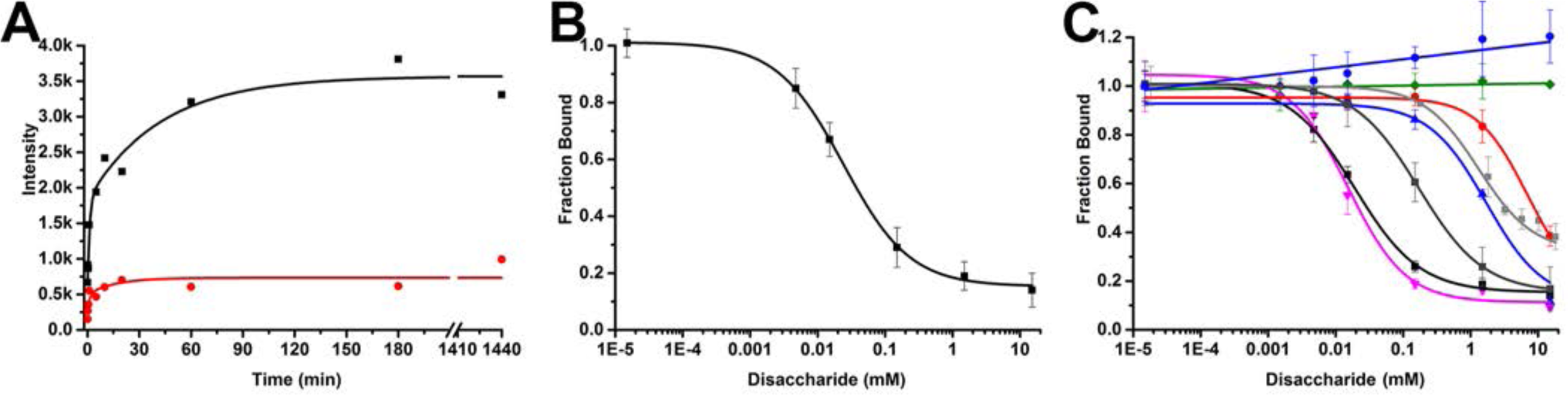
Interaction between PCV2 and heparin conjugated chromatography sorbent. A) Binding kinetics between PCV2 VLP and the chromatography sorbent: 370nM (black) and 92nM (red) concentration of VLP. B) Elution of PCV2 from the chromatography sorbent in the presence of increasing concentration of heparin. C) Competition assays testing the ability of various GAGs and polysaccharides to displace PCV2 from the chromatography sorbent: (red) dp6 heparin (6-hexose), (blue) dp10 heparin (10-hexose), (dark grey) dp20 heparin (20-hexose), (black) dp36 heparin (36-hexose), (magenta) dextran sulfate 8kDa, (brown) dextran sulfate 6K, (light grey) chondroitin sulfate B 41kDa, and (green) hyaluronic acid. Error bars represent the standard deviation of three measurements. Images were made with Origin 2016 (OriginLab).

To determine the chemical moieties of GAGs that PCV2 recognizes for cellular attachment, we employed a competition assay to identify the effectiveness of different ligands to compete with the mentioned chromatography sorbent for binding PCV2 (**Figure 2C**). We examined the length of heparin, the pyranose structure, sulfate content, and branched polysaccharide chains. The results demonstrate that while the efficiency of heparin dp6 (6-hexoses) (**Figure 2C, S2C**), dp10 (10-hexoses) (**Figure 2C, S2D**) and dp20 (20-hexoses) (**Figure 2C, S2E**) to inhibit 15kDa heparin from binding to PCV2 increases, they are not nearly as effective as dp36 (36-hexoses, 12kDa) heparin (**Figure 2C, S2F**). A plausible explanation could be that there are multiple weak binding sites on the PCV2 capsid, and that longer oligosaccharides have higher affinity to PCV2 because they can bind to multiple sites concurrently. It has been shown that heparin fragments below dp18 are helical, extended and rigid but become more bent with increasing size (37). Thus the dp6 and dp10 may bind to a few binding sites on a PCV2 facet, while the dp20 and dp36 heparin may bend and bind to multiple facets on PCV2.

To address if a polysaccharide with high sulfate content can interact with PCV2, we examined the ability of dextran sulfate (8kDa) to inhibit the interaction between PCV2 and the chromatography sorbent (**Figure S2G**). The dextran sulfate (8kDa) used in our studies is a highly branched polysaccharide, with glucose repeating units connected via 1→4 linkages and branching points at 1→6. There are 2.8 sulfates per glucose monomer for this product (personal communication with Alfa Aesar manufacturer). Dextran sulfate is the ligand most efficient at inhibiting the interaction between PCV2 and 15kDa heparin (**Figure 2C**). However dextran, which has the same-branched structure as dextran sulfate, appears to slightly enhance the affinity of the VLP for the sorbent (**Figure S2H**). This may be due to dextran acting as a crowding agent (38). The ability of dextran sulfate to inhibit the PCV2-heparin interaction while dextran slightly enhances the interaction suggests that the sulfates are sufficient for binding to PCV2, and that the backbone polysaccharide common to dextran and dextran sulfate does not appear to interact with PCV2.

To address if longer GAGs with lower sulfate content exhibit reduced affinity for PCV2, we examined the ability of 1.6MDa hyaluronic acid (**HA**) and 41kDa chondroitin sulfate B (**CSB**) glycosaminoglycans to inhibit the interaction between PCV2 and 15kDa heparin. HA demonstrates no affinity for PCV2 (**Figure 2C, S2I**). HA possesses a single carboxylate on each of its repeating disaccharide units. The hydrogen bonds formed within and between its repeating disaccharides produce stiff helical polysaccharide chains that entangle at very low concentrations (39). Thus, the carboxylate, stiff, and entangled properties of HA may not provide the appropriate interface for PCV2 to bind. The CSB used in our studies is longer and more flexible (40) than the 12kDa heparin we used; however, CSB demonstrates a much lower affinity for PCV2 than heparin. This may be because of the lower sulfate content of CSB (an average of 1.1 sulfate per repeating disaccharide) when compared to heparin (an average of 2.4 sulfates per repeating disaccharide) (**Figure 2C, S2J**).

Collectively, our data indicate that the PCV2-heparin interaction is reversible, the strength of the interaction increases with longer lengths of heparin, oligosaccharides high in sulfate content bind with greater affinity, long oligosaccharides with low sulfate content bind weakly, and that the interaction between PCV2 and the polysaccharide is predominantly driven by sulfates on the polysaccharide.

### A capsid subunit can bind heparin via multiple sites

To identify the binding sites of heparin on PCV2 we determined the cryo-EM image reconstruction of PCV2 in the absence and presence of 12kDa (dp36) porcine intestinal mucosa heparin at pH 7. The purpose of the unliganded image reconstruction was to identify if a conformational change occurred in the subunit upon binding to heparin. We generated icosahedral image reconstruction of the unliganded and heparin-liganded PCV2 to 3.3Å and 2.8Å resolution, respectively. The image reconstructions are of suitable quality to allow us to confidently model the atomic coordinates of PCV2 (**Table 1, Figure 3A**). The refined atomic models for the two image reconstructions overlay with a Cα RMSD of 0.27Å, and an all atom RMSD of 0.98Å (41). These differences are small and can be a result of modeling error at these resolutions (**Figure 3B**) (42, 43). The symmetrized heparin-liganded image reconstruction does not allow us to confidently identify the molecular envelopes for heparin.

**Table 1:**
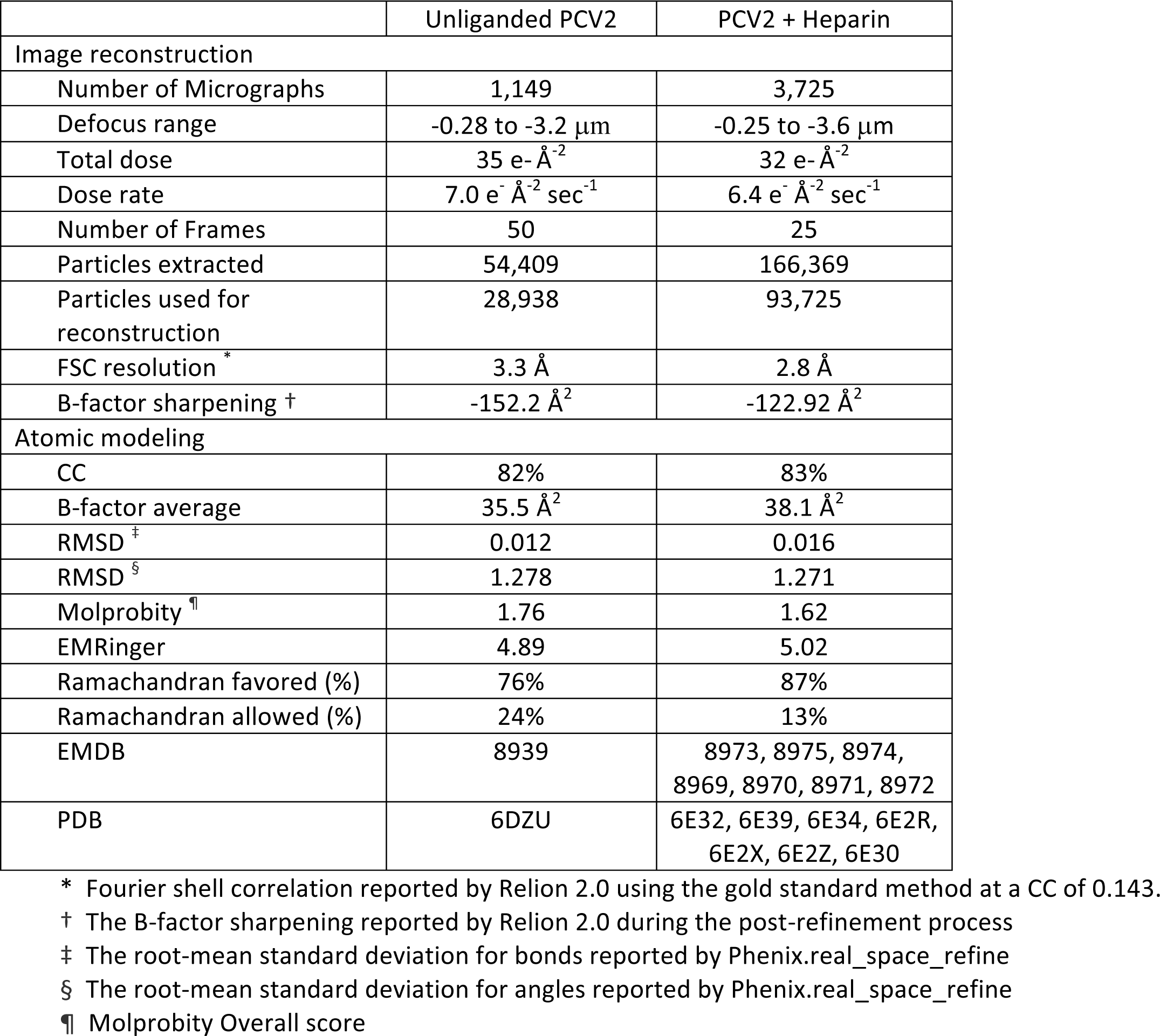
Statistics of cryo-EM image reconstructions and refined atomic coordinates.

**Figure 3.**
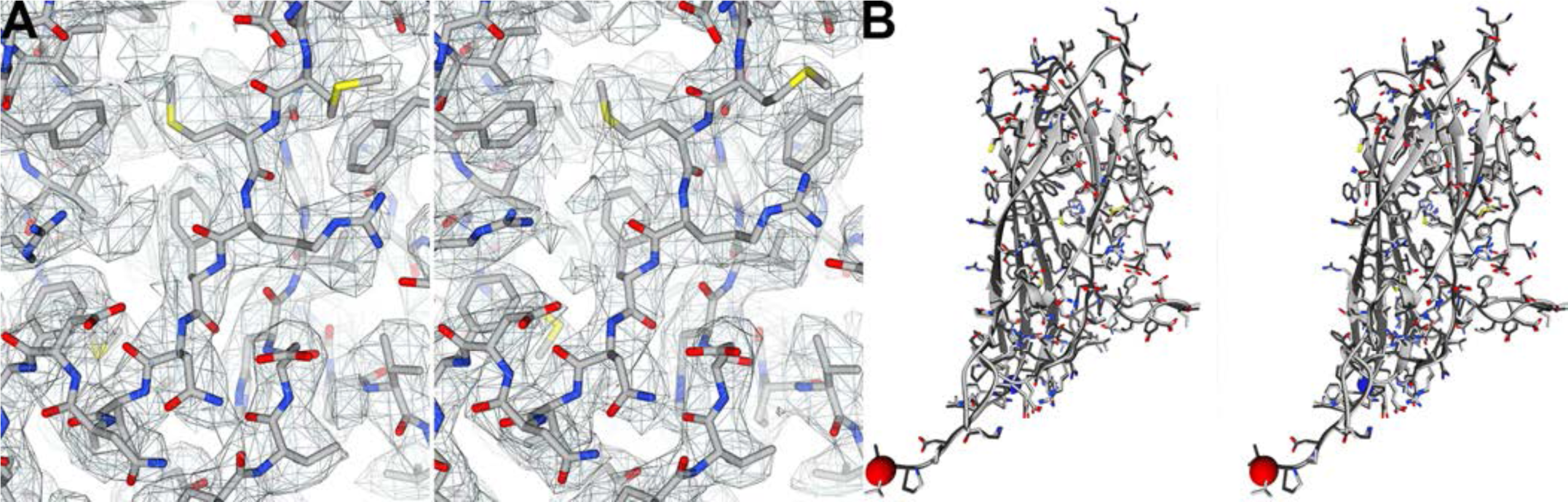
Symmetrized cryo-EM image reconstructions of unliganded and heparin liganded PCV2. A) Full atom models of an unliganded (left) and heparin liganded (right) PCV2 subunits shown as stick models. Overlaid onto the stick models are mesh representations of the corresponding 3.3Å (left) and 2.8Å (right) symmetrized image reconstructions. The unliganded image reconstruction is rendered at 1.5-sigma, and the liganded image reconstruction is rendered at 2.6-sigma. The rendering thresholds correspond to 100% mass content –see methods and materials section. B) Cross eye stereo representation comparing the atomic models derived from the unliganded (light grey) and liganded (dark grey) image reconstructions. The backbone atoms are depicted as a ribbon diagram, and the side chains are shown as stick models. Blue and red spheres represent the N- and C-termini, respectively. Images generated with UCSF Chimera (64).

We postulated that if each subunit possessed multiple binding sites for heparin but only one site could be occupied per subunit then each capsid might be composed of a unique combination of subunits liganded to heparin; thus, icosahedral-averaging strategy could eliminate the molecular envelope of heparin. To test this possibility, we employed the symmetry expand capability of Relion on the alignment parameters from the image reconstruction where icosahedral symmetry (I1) was imposed (44). Briefly, the symmetry expansion protocol generates a copy of each particle for each symmetry operator used during the symmetrized image reconstruction. In our case, 60 equivalent particles were generated for each imaged VLP because 60 matrices describe the I1 symmetry used by Relion. The orientation parameters of these particles are determined by modifying the orientation parameters of the parent particle by the 60 symmetry operators that describe the I1 symmetry. Consequently, a single imaged VLP that was assigned an orientation parameter for I1 symmetry is replaced with 60 copies of the VLP assigned orientation parameters for C1 symmetry. These 60 orientation parameters are related to one another via I1 symmetry. The consequence of this strategy is to align the projections of all the subunits from the VLP to one another (45). We then used Frealign to perform focused classification of a subunit (46, 47). Seven of the ten classes exhibit molecular envelopes distinct from the symmetrized image reconstruction (**Figure 4**). To generate a difference map for each class average we subtracted from each class average an image reconstruction generated using the same number of particles as the class average –these particles were randomly selected from the symmetry expanded data set. Calculation of local resolution for each of the ten classes, with MonoRes, suggests that the resolution of the peaks vary from 7Å to 15Å (**Figure S3**) (48). The difference peaks were low-pass filtered to 7Å resolution and rendered at 27-sigma (**Figure 4C**). We used 27-sigma as the criteria to remove small and spurious density.

**Figure 4.**
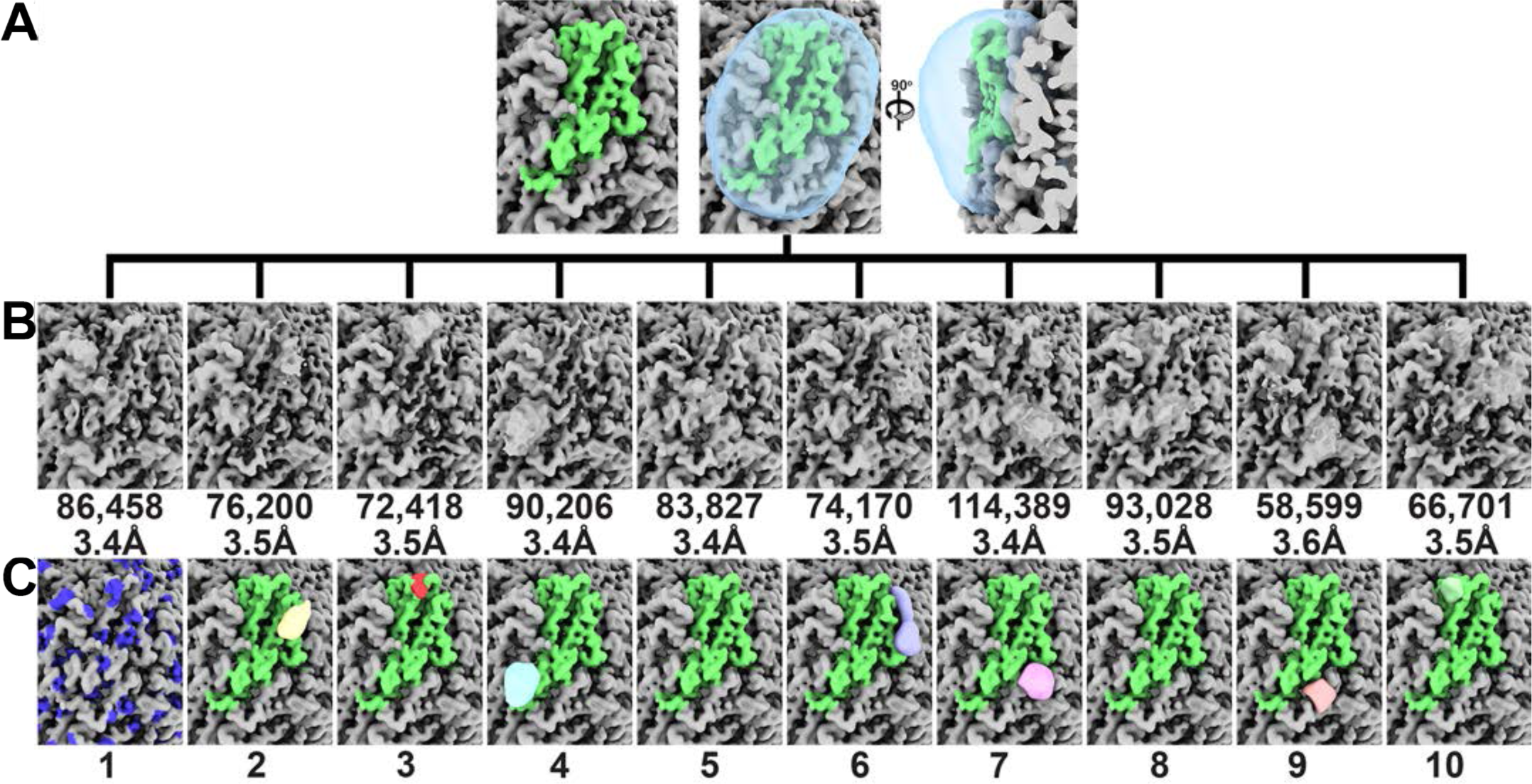
Focused classification of the PCV2 VLP liganded to heparin image reconstruction. Symmetry expansion of 13,600 particles generated 816,000 capsid subunits for analysis. The image reconstructions are shown at 2.6-sigma. A) The molecular envelope for a subunit is shown in green. The mask used for the classification is shown as a transparent cyan blob. B) Ten classes were requested. The number of particles and resolution reported by Frealign is written below each class. C) Difference peaks for each class average shown in B. There is no difference peak for Class 1. The blue patches represent the molecular envelope of Arg/Lys side chains. Class numbers referred to in the text are written at the bottom. Images made with UCSF ChimeraX (69).

The difference peaks may correspond to segments of the 12kDa heparin molecule incubated with PCV2. We note that the amorphous shapes of these peaks indicate that heparin may use multiple binding modes/poses to bind to the capsid. Indeed, heparin is known to use multiple binding modes/poses to interact with proteins (28). We modeled segments of heparin as rigid bodies into the difference peaks to see if the models were chemically reasonable. We then examined if mutation of the basic and polar amino acids that interact with the sulfates in our models diminished the mutant VLP’s affinity for heparin. The VLP for these studies were generated using an *in vitro* assembly protocol of capsid protein expressed in *E. coli* –see Materials and Methods.

Class 1 (11% of subunits) does not possess a difference peak and thus represents an unliganded subunit. The difference peak for class 2 (9% of subunits) is located near Arg73 and Lys132 of a subunit. A four-hexose segment of heparin can be modeled into this difference peak (**Figure 5A**). Mutation of Arg73 to Ala diminishes the capsid-heparin affinity to 68% of wild type, whereas mutation of Lys132 to Ala abolishes capsid assembly (**Figure 5F**). The difference peak for class 3 (9% of subunits) is located near Lys58 and Arg59, but is too small to accommodate heparin. The difference peak for class 4 (11% of subunits) is near Arg89, and next to the icosahedral 2-fold axis of symmetry. A three-hexose segment of heparin can be modeled into this density (**Figure 5B**). Mutation of Arg89 to Ala diminishes the capsid-heparin affinity to 46% of wild type (**Figure 5F**). The difference peak for class 5 (10% of subunits) is extremely weak. The difference peak for class 6 (9% of subunits) runs between adjacent facets. It is located near Arg186 and Lys227 of the adjacent subunit. A five-hexose segment of heparin can be modeled into this difference peak. A number of hydrogen bonds are also possible between the sulfates of heparin and the side chains of Thr131, Tyr138, and Ser86 (**Figure 5C**). Mutation of Arg186 to Ala diminishes the capsid-heparin affinity to 78% of wild type, whereas mutation of Lys227 to Ala abolishes capsid assembly (**Figure 5F**). The difference peak for class 7 (14% of subunits) is distal to Arg/Lys, but a five-hexose segment of heparin modeled into the difference peak suggests that hydrogen bonds can form between the sulfates of heparin and the side chains of Asn128, Gln188, and the three Tyr173 at the icosahedral 3-fold axis (**Figure 5D**). Mutation of Asn128 or Gln188 to Ala diminishes the capsid-heparin affinity to 85% and 84% of wild type, respectively (**Figure 5F**). The difference peak for class 8 (12% of subunits) is also very weak. The difference peak for class 9 (7% of subunits) is located near Arg73 and Lys132 of the neighboring subunit, and is equivalent to the peak of class 2. This class has been identified by our procedure because the mask used for classification includes regions of neighboring subunits that are related to one another via the icosahedral symmetry of the capsid. Given that this is a redundant class, we exclude it from further analysis. The difference peak for class 10 (8% of subunits) is located near the icosahedral 5-fold axis, and surrounded by Lys58 and Arg59 of one subunit and Lys63 of a neighboring subunit. A sulfate was observed at this location in the crystal structure of PCV2 (35). A two-hexose segment of heparin can be modeled into this density (**Figure 5E**). Mutation of either Lys58 or Arg59 to Ala diminishes PCV2-heparin interaction to 69%, while mutation of Lys63 diminishes the affinity to 38% (**Figure 5F**).

**Figure 5.**
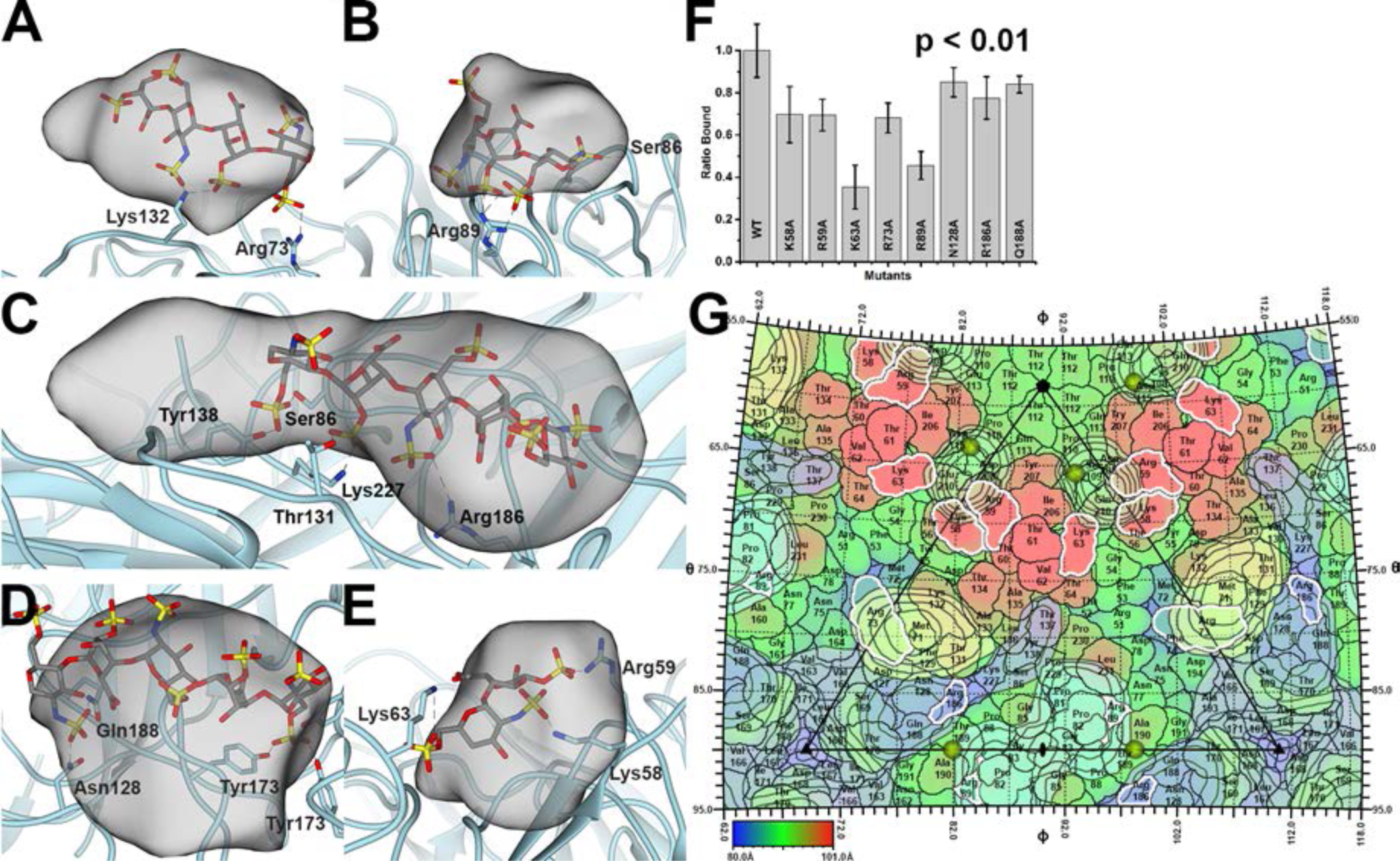
PCV2 heparin interaction. The PCV2 capsid is shown as a cyan ribbon cartoon. The difference peak (at 27-sigma) is shown as a grey transparent blob –low pass filtered to 7Å. A modeled heparin is shown as a grey stick model. The dashed lines represent hydrogen bonds and/or salt bridges to the sulfates (red and yellow sticks) of heparin. A) Class 2 binding site. B) Class 4 binding site. C) Class 6 binding site. D) Class 7 binding site. E) Class 10 binding site. Images made with UCSF Chimera (64). F) Bar graph representing the efficiency of 370nM mutant PCV2 VLP for binding to heparin, with wild-type (WT) VLP having been normalized to 1. The error bars represent errors from five independent measurements. Image made with Origin 2016 (OriginLab). The p-value was calculated using the *ranksum* (Wilcoxon rank sum test) routine of MATLAB 2018a (MathWorks). G) The surface amino acids of PCV2 are plotted on a stereographic projection and colored from blue (80Å) to red (101Å) according to the distance from the center of the particle –see color bar. Overlaid are contour plots at 27-sigma (black outline) for the six difference peaks. The plots are colored according to Figure 4. Residues with a white outline were mutated to Ala in this study, assembled into capsids, and tested for their ability to bind to heparin. Note that Lys137 and Lys227 have not been colored because they did not assemble into capsids and a binding assay could not be performed. The sulfates observed in the crystal structure of PCV2 are shown as gold spheres (35). The icosahedral 5-fold, 3-fold, and 2-fold symmetry elements are depicted as a black pentagon, triangle, and ellipse respectively. Figure generated with RIVEM (49).

To compare the difference peaks on the surface of the PCV2 capsid, we generated a stereographic projection of the PCV2 surface with the program RIVEM (**Figure 5G**) (49). Overlaid onto this image are contour plots of the symmetrized difference peaks, the sulfates observed in the PCV2 VLP crystal structure, and the 532-icosahedral symmetry elements (35). The difference peaks have been symmetrized to visualize all the binding sites available on the surface of the capsid. The sulfates observed in the crystal structure are located near the difference peaks from classes 4 and 10. The stereographic projection demonstrates that the differences peaks from classes 6 and 7 partially overlap and share a number of amino acids that include Asn128 and Gln188.

The resolution of the image reconstructions from each class is of sufficient quality to refine the atomic models. Refining the atomic model generated from the image reconstruction with imposed icosahedral symmetry into the subunit from each of the seven classes does not identify significant movements in the main chain atoms that could be associated with a conformational change upon binding to heparin. The largest Cα RMSD between subunits from two classes is 0.313Å.

### The occupied heparin binding sites do not follow the icosahedral symmetry of the capsid

The result of focused classification is to assign each of the sixty subunits belonging to an imaged VLP to one of ten classes. We used this information along with the inverse of the Relion I1 matrices to position the difference peaks onto the originating VLPs (**Figure 6**). We refer to these models as composite VLP, as they are composed of a symmetrized PCV2 image reconstruction and the positioned difference peaks. The number of difference peaks present per VLP follows a Gaussian distribution: with VLP possessing a minimum of 18 peaks (**Figure 6A, S5**), an average of 31 peaks (**Figure 6B, S5**), and maximum of 43 peaks (**Figure 6C, S5**). The dp36 heparin used to generate this data prompted us to look for occupied binding sites that could accommodate its physicochemical properties (**Figure 6**). Indeed, we can manually model dp24 and dp36 heparin (PDB entries 3IRJ and 3IRL) as rigid bodies into these composite VLPs (**Figure 6A, 6B, 6C**). The bend in the dp36 heparin allows it to bind binding sites distributed across two adjacent facets of the capsid.

**Figure 6.**
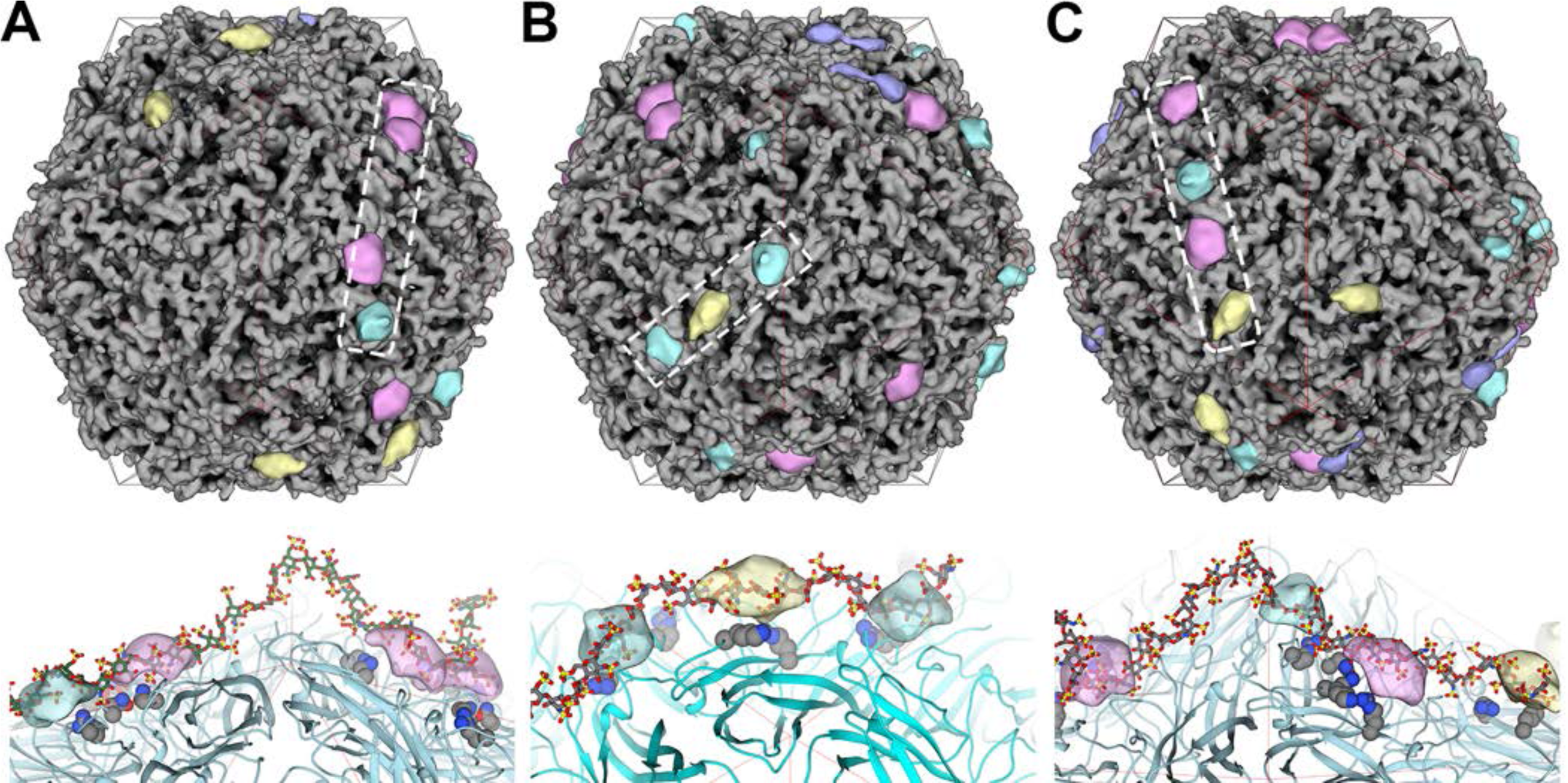
Composite VLPs, where each model represents an imaged VLP-heparin complex. The coloring scheme used for the difference peaks is identical to Figure 4. Top, the dashed box outlines the occupied binding sites that a heparin oligosaccharide is modeled into (bottom). Bottom, a heparin oligosaccharide (grey stick model) occupying the difference peaks (transparent blobs) identified by the dashed box (top). The binding site residues are shown as grey space filling models and the PCV2 capsid is shown as a cyan ribbon model. A red icosahedral cage distinguishes the facets. A) VLP with 18 binding sites occupied by heparin. A dp36 heparin has been modeled into the outlined sites. B) VLP with 31 binding sites occupied by heparin. A dp24 heparin has been modeled into the outlined sites. C) VLP with 40 binding sites occupied by heparin. A dp36 heparin has been modeled into the outlined sites. Images made with UCSF Chimera (64).

## Discussion

The cellular attachment factors responsible for initiating infection of porcine monocytic 3D4/31 and porcine kidney epithelial PK-15 cells by PCV2 were demonstrated to be heparan sulfate (**HS**) and chondroitin sulfate B (**CSB**) using competition assays, enzymatic removal of HS and CSB, and mutant CHO cells deficient in HS (*pgs*D-677) or HS and CSB (*pgs*A-745) (25). Both HS and CSB are long linear polysaccharides known as glycosaminoglycans (**GAGs**). HS is composed of two alternating domains: a variably sulfated and rigid NS domain and a non-sulfated and flexible NA domain (26, 29, 50). Three to eight repeating disaccharides generate the NS domain, and two to twelve repeating disaccharides generate the NA domain. The NS domain has a chemical structure similar to heparin with a length of dp6 (6-hexoses) to dp16 (16-hexoses) (**Figure S1)**. The NS domains bind to a plethora of ligands that include, but are not limited to, cytokines, enzymes and their inhibitors, receptors, chemokines, and pathogens (26). A number of non-enveloped icosahedral viruses are known to use HS for cellular attachment. Examples include serotypes of adenovirus (51), serotypes of adeno-associated virus (52), foot-and-mouth disease virus (53), and human papillomavirus (54). Here we note that while heparin is routinely used as an analogue of HS (29), it is not a cellular attachment factor. Heparin is stored in the secretory granules of mast cells and released into the vascular system as an anti-inflammatory agent (30). An extensive review by Xu and Esko describes that a large portion of the binding energy between heparin and proteins can be attributed to the ionic interaction between the Arg/Lys amino acids of proteins and the sulfates of heparin. However, polar amino acids such as Asn, Gln and His can also make hydrogen bonds with the sulfates of heparin, and can contribute 20-70% of the binding energy. Analysis of structures deposited into the PDB indicates that heparin binding sites vary in size (200Å^2^ to 360Å^2^), number of lysine and arginine residues (0 to 7), and number of polar amino acids (0 to 5). Interestingly, the size of the interface and number of basic residues in the binding site do not predict the affinity of the interaction. Protein-heparin binding strength also vary from 0.3nM to 7μM (K_d_). The majority of the heparin binding amino acids are present in the loops of the proteins (28).

We have used an *in vitro* competition assay to demonstrate that PCV2 virus like particles (**VLP**) bind to increasing lengths of heparin fragments with greater affinity. The binding trend suggests that the PCV2 VLP may possess multiple weak binding sites such that longer fragments of heparin are able to bind additional sites for producing greater affinity –a phenomenon referred to as avidity (55). Our assays also demonstrate that PCV2 has a greater affinity towards dp36 heparin (12kDa) than the much longer 41kDa CSB. This is in agreement with the cellular competition assays that demonstrate the incubation of PCV2 with heparin reduces infection greater than incubation of PCV2 with CSB (25). Moreover, infection of Chinese hamster ovary (**CHO**) mutant cell lines *pgs*A-677 (lacking HS) was shown to be reduced to ^∼^28% that of wild-type, while infection of the mutant cell lines *pgs*A-745 (lacking HS and CSB) was reduced to ^∼^10% that of wild-type. Suggesting that HS may be the preferred attachment factor on the surface of the cell.

Our *in vitro* competition assays also demonstrate that PCV2 VLP exhibit greater affinity towards sulfated polysaccharides. In particular, the observation that PCV2 VLP binds to dextran sulfate (**DS**) 8K but not to dextran 6K suggest that PCV2 recognizes and interacts with the sulfates of the polysaccharide. Indeed, our competition experiments demonstrate that DS is the most potent of competitors tested in our assays, suggesting that DS may be used as an inhibitor of PCV2 infection.

Our structural data identify six class averages where additional molecular envelopes modify the molecular envelope of the capsid subunits. Subtracting the molecular envelope of an unmodified capsid from these class averages generates six difference peaks. The difference peaks are large enough to accommodate heparin oligosaccharides. Heparin oligosaccharides can be modeled into the difference peaks such that their sulfates are in close proximity to amino acids Lys58, Arg59, Lys63, Arg73, Arg89, Lys132, Asn128, Arg186, Gln188, and Lys227. Single point substitution of these amino acids to Ala diminish the capsid’s affinity for heparin, and thus provide biochemical support that the difference peaks represent bound heparin. A single mutation is not sufficient to tremendously diminish binding of heparin; therefore, we anticipate that multiple mutations may be necessary to eliminate the interaction. Comparison of the difference peaks suggests that the peaks belonging to two class averages are equivalent. Thus heparin can bind to one of five binding sites per PCV2 capsid subunit, such that a PCV2 VLP can possess a maximum of 60 sites occupied by heparin. Given that each difference peak originates from a subunit at a specific position in a VLP, the difference peak can be mapped to that position in the VLP to generate a heparin-liganded molecular envelope for each imaged VLP. These molecular envelopes demonstrate that heparin asymmetrically decorates the surface of the capsid rather than the symmetric decoration demonstrated for a number of icosahedral viruses. This may be because prior structural studies of icosahedral viruses in complex with heparin utilized averaging techniques that may have eliminated the asymmetric distribution of heparin on the capsid surface (52, 53, 56–59). The packing of capsids into a crystal for X-ray crystallography inherently imposes the averaging of structures (60). Similarly, cryo-EM image reconstructions with imposed icosahedral symmetry average non-icosahedral entities to symmetric structures with reduced occupancy or amorphous blobs (e.g. the molecular envelope of vial genomes). However, the use of symmetry expansion and focused classification forgo symmetry averaging to provide a more detailed analysis.

Our modeling suggests that the reported solution structures of dp24 and dp36 heparin (37) can bind to adjacent facets of the PCV2 capsid. This is primarily because of the bent structure of these oligosaccharides (37). Consequently, it is anticipated that shorter heparin lacking such a bend will be unable to interact with adjacent facets. The longest NS domain of HS is reported to be eight disaccharides (dp16) (28) and thus unlikely to bend and bind to adjacent facets. This creates the likelihood that an NS domain binds to a single facet and the neighboring flexible NA domain allows the HS chain to bend for the subsequent NS domain to bind to an additional facet on the surface of PCV2. Heparan sulfate possesses 40-300 sugar molecules (20-150nm long), and is composed of multiple alternating NS (dp6-dp16) and NA domains (dp4-dp24) (26, 28, 50).

In conclusion, we demonstrate that PCV2 capsids bind increasing lengths of heparin with greater affinity, and that this interaction is predominantly driven by the affinity for the sulfates than the pyranose. Our structural studies visualize heparin to bind one of five binding sites per subunit, and the binding of heparin to the capsid to not adhere to the icosahedral symmetry of the capsid. Binding of heparin does not induce a conformational change in the capsid. We also show that longer heparin oligosaccharides (dp24 and dp36) can be modeled to bind adjacent icosahedral facets. Our studies strongly imply that polymers of sufficient length and sulfate content, including heparin 12kDa and dextran sulfate 8kDa, may act as inhibitors of PCV2 infection, and that such inhibitors may act as therapeutics in swine farms where PCV2 infection is present. Future studies include assessing if capsid dynamics are altered upon binding of heparin, if certain binding sites are dispensable for binding to heparin and heparan sulfate, and if the asymmetric distribution of heparin on the capsid surface is important for the life-cycle of the virus.

## Materials and Methods

### PCV2 VLP expression and purification

The codon sequence for the PCV2 capsid protein (GeneBank ACN59889.1) was optimized by GenScript for Baculovirus expression system. GenScript also generated the Baculovirus stock used in these studies. *Trichoplusia ni* insect cells were propagated in ESF921 medium (Expression System) and infected with the Baculovirus at a multiplicity of infection (MOI) of 0.1. Infected cells were grown for 5 days (50% viability according to Trypan blue staining), and harvested via centrifugation at 3,000xg for 20 minutes.

PCV2 VLP was purified from the nucleus of the infected cells. Cells from 500ml of media were gently centrifuged at 800xg for 10 minutes at 4°C. The cell pellet was suspended and lysed in 20mM 4-(2-hydroxyethyl)-1-piperazineethanesulfonic acid (HEPES) pH 7.5, 250mM Sodium Chloride (NaCl), 2mM Magnesium Chloride (MgCl_2_), 1mM Tris(2-carboxyethyl)phosphine hydrochloride (TCEP), 1% (v/v) NP-40, and placed on a revolver at 4°C for 30 minutes. The lysate was centrifuged for 30 minutes, 720xg, at 4°C to pellet the nuclei. The nuclei pellet was suspended in nuclear lysis buffer: 20mM HEPES pH 7.5, 500mM NaCl, 2mM MgCl_2_, 0.2mM TCEP, 2mM Deoxycholate, 1μl salt activated nuclease (Sigma-Aldrich, SRE0015-5KU) and lysed using a sonicator. The lysate was centrifuged at 28,000xg for 30 minutes at 4°C. The supernatant was overlaid on a 30% sucrose cushion and centrifuged at 184,000xg, 11°C, 2.5 hours using a 45Ti Beckman Coulter rotor. The pellet was suspended in 2ml of nucleus lysis buffer and processed on a discontinuous 20 to 70% sucrose gradient. The gradient was centrifuged using a SW40 Beckman Coulter rotor at 160,000xg, 4°C, and 18 hours. The fraction containing the VLP was visually identified using light scattering and SDS-PAGE. The fraction containing VLP was dialyzed overnight in 20mM HEPES pH 7.0, 250mM NaCl, 0.2mM TCEP, and 2mM ethylenediaminetetraacetate (EDTA). Samples were concentrated using a 100kDa MWCO ultrafiltration device (Pall Corporation), and stored at 4°C. The sample concentration was estimated to be 370nM using SDS-PAGE with Coomassie staining against a standard of similar molecular weight with a known concentration.

### Binding studies

Heparin HyperD™ M Affinity Chromatography Sorbent conjugated to 15kDa porcine intestinal mucosa heparin was purchased from Pall Corporation. HyperD™ M was washed and equilibrated with 20mM HEPES pH 7.0, 250mM NaCl, 0.1mM TCEP, 2mM EDTA and allowed to settle to the bottom of the tube. Each reaction contained 2µl of settled HyperD™ M and the indicated amount of VLP (see below). Samples were incubated at 20°C for the indicated amount of time (see below –kinetic binding assays) on a revolver to allow PCV2 to bind to the sorbent. The HyperD™ M sorbent was gently pelleted at 100xg for 30s, washed two-times using the equilibration buffer, suspended in water, and processed using SDS-PAGE. The PAGE was stained using InstantBlue™ from Expedeon, and band intensities were quantitated using the Li-Cor Odyssey Blot Imager equipped with Image Studio™ Software version 5.0. The reported Fraction Bound is the quotient of the intensity for each band (**Figure S2**) and the intensity of the band for the lowest concentration of polysaccharide tested. The washes did not remove VLP from the sorbent, as exhibited by SDS-PAGE (data not shown).

Kinetic binding assays: 4µl of VLP at concentrations of 370nM and 92.5nM were incubated with 2µl of HyperD™ M for 5 seconds (**s**), 10s, 30s, 1 minute (**m**), 5m, 10m, 20m, 1 hour (**h**), 3h and 24h on a revolver at 20°C. The sorbent was processed as described above.

#### Elution assays

VLP bound to HyperD™ M sorbent (as described above) was incubated with increasing concentrations of porcine intestinal mucosa heparin (see below –competition assays) for 3h at 20°C. The bound VLP was quantitated as described above.

#### Competition assays

PCV2 was incubated with the GAG reagents at 20°C for 3h prior to incubation with 2μl of settled HyperD™ M sorbent for 3h. The sorbent was processed as described above for SDS-PAGE.

### Glycosaminoglycans and polysaccharide source

Heparin oligosaccharides (HO06, HO10, HO20), and chondroitin sulfate B (DS01) were purchased from Galen laboratories, and dissolved in the VLP dialysis buffer. Porcine intestinal mucosa heparin (average molecular weight 12kDa) was purchased from JT Baker, hyaluronic acid and dextran sulfate 8kDa from Alfa Aesar, and Dextran 6000 (Sigma-Aldrich). All reagents were used within 48 hours after being dissolved in the VLP dialysis buffer. Concentrations were calculated based on the disaccharide molecular weight of 659 g/mol for heparin, 551 g/mol for chondroitin sulfate B, 389 g/mol for Hyaluronic acid, and 728 g/mol for dextran sulfate 8kDa. Samples were stored at 4°C.

### Electron microscopy

Ultrathin carbon film on Lacey Carbon support film (TedPella, 01824) were used for cryo-electron microscopy. Grids were glow discharged using a PELCO easiGlow™ for 30 seconds. Unliganded and heparin liganded VLP (4µl) were adsorbed to the grid for 15 seconds, blotted with a force of 2 for 6 seconds, and plunge frozen into liquid ethane using an FEI Mark IV vitrobot. Data for the unliganded PCV2 was collected using a side entry FEI Titan Halo with the Leginon software (61). Images were collected at a calibrated magnification of 105,000x on a Gatan K2 camera operating in counting mode. Cryo-EM grids for the PCV2 liganded to heparin were prepared by incubating 20µl of the VLP with 7µl of 60mg/ml 12 kDa heparin for 3 hours. The mixture was then incubated with 2µl of HyperD™ M sorbent for 3 hours at 20°C on a revolver to remove VLP that was not bound to heparin. Cryo-EM grids were prepared as described for the unliganded PCV2. Data for the heparin liganded PCV2 was collected on a Titan Krios with the Leginon software. Images were collected at a calibrated magnification of 105,000x on a Gatan K2 camera that follows a GIF Quantum Energy Filter with a slit width of 15eV. Data collection parameters for both data sets are reported in **Table 1**.

### Image reconstruction

Motion correction was performed using the MotionCor2 package (62). The first frame of each movie was discarded during the alignment. With the exception of a patch 5 option, the default options were used for the alignment and dose weighting. Particles were automatically detected using Gautomatch v0.53 and per-particle contrast transfer function (**CTF**) estimation was performed on the aligned micrographs using Gctf v0.50 (63). Particles were extracted from the dose-weighted micrographs using the coordinates identified by Gautomatch. Particles from the unliganded dataset were extracted into boxes of 256×256 pixels, and particles from the heparin-liganded data set were extracted into boxes of 300×300 pixels. Three iterations of reference free 2D classification with a diameter of 250Å, 128 classes, and default Relion 2.0 parameters were used to discard bad particles between each iteration (44). The PCV2 crystal structure (PDB entry 3R0R) was used to generate a 60Å resolution initial model using the *molmap* function of UCSF Chimera (64). 3D classification was carried out with the default Relion 2.0 parameters using C1 symmetry. A total of 39,048 particles for the unliganded dataset and a total of 118,853 particles for the heparin-liganded dataset were classified using 3D classification. A total of 4 classes were requested for the unliganded dataset and a total of 10 classes were requested for the liganded dataset. From each dataset, particles from the 3D class with the highest resolution image reconstruction were selected for further full resolution refinement using I1 symmetry. The default parameters of Refine3D in Relion 2.0 were used to achieve the highest resolution possible. For each data set, binary masks for postprocessing were generated using the *relion_mask_create* program of Relion 2.0 using the following options: 1) a low pass filtered final image reconstruction at 15Å resolution, 2) a threshold value just high enough to eliminate noise exterior to the PCV2 capsid, 3) mask extension by 7 pixels, and 4) additional 2 soft edge pixels expansion. Masks were manually inspected with UCSF Chimera to ensure that no cavities existed in the interior of the masks. Postprocessing was performed with the default parameters of Relion 2.0 (44).

The particle orientation parameters attained from the full resolution refinement described above were expanded from I1 to C1 symmetry using the *relion_particle_symmetry_expand* program of Relion 2.0 for 13,600 heparin-liganded particles. The consequence of this is to generate 60×13,600 (816,000) subunits that have been aligned to one another. The molecular envelope for a PCV2 subunit that was refined (see below) into the I1 image reconstruction was calculated to 25Å resolution using the *molmap* function of UCSF Chimera, threshold masked, extended by 7 pixels, and extended by 2 soft edge pixels using the *relion_mask_create* program of Relion 2.0 (44, 64). The mask was manually moved radially outward to include area peripheral to the capsid in case heparin may be present on the surface of the capsid. The alignment parameters and images were converted to a format for Frealign using the *relion_preprocess* function of Relion and the conversion script provided by the Frealign authors. Twenty rounds of masked classification (10 classes), no alignment, were performed with Frealign (46, 65). Difference maps were calculated using the *vop* function of UCSF Chimera (64). The image reconstruction from each class was subtracted from an image reconstruction generated using the same number of particles randomly selected from the data set. This was done so as both image reconstructions have comparable signal-to-noise ratio. The image reconstructions from the randomly selected particles were visually indistinguishable from the icosahedrally averaged image reconstruction.

The image reconstructions were rendered at 100% mass content of the capsid shell (60×27,853Da, or 1.67 MDa). The threshold level to render the images was calculated using the *volume* command from EMAN with the pixel size of the image reconstruction (66).

### Structure refinement

The reported crystal structure of PCV2 (35) was manually docked into the symmetrized image reconstructions using UCSF Chimera. The resulting coordinates were iteratively refined using *phenix.real_space_refine* from the Phenix software package with non crystallographic symmetry (**NCS**) constraints applied, and manual fitting with Coot (67, 68). Heparin molecules (PDB entries 3IRI, 3IRJ, 3IRK, and 3IRL) were treated as rigid bodies and manually fitted into the difference peaks of the five classes described above. The capsid subunits pertaining to these molecular envelopes were further refined using manual fitting with Coot and *phenix.real_space_refine –* without NCS.

### Generating Composite VLP, representative models of individual VLP

The focused classification identifies one of ten class averages that represents each subunit of a symmetry expanded VLP, and the difference peaks represent each of these ten classes. Consequently, the class averages can be combined with the symmetry elements used in the symmetry expansion process to generate a model of each imaged VLP. The icosahedral image reconstructions were generated using the I1 symmetry of Relion, and symmetry expansion was performed using the I1 symmetry matrices. Consequently, inverse matrices for the Relion I1 matrices should position the class averages, or their corresponding difference peaks, onto the VLP from where they originate. Inverse matrices were calculated for the I1 symmetry matrices using MATLAB R2018a (MathWorks). These symmetry operators were then applied to difference peaks representing each subunit of a VLP with UCSF Chimera (64).

### Mutagenesis and binding efficiency

Single point mutants of the PCV2 capsid were made to the consensus sequence construct described in Khayat et al. 2011 (35). Briefly, this construct possesses a hexahistidine and a thrombin cleavage site at the N-terminus in lieu of the forty-two amino acids in the GeneBank ACN59889.1. The remaining amino acid sequence is identical. Mutations were made using the New England Biolabs Inc. site directed mutagenesis kit. Sequences were confirmed using the services of Genewiz. Proteins were purified as previously described (35). Virus like particles (**VLP**s) were assembled by mixing equal volumes of the capsid protein at 37µM concentration with 12% polyethylene glycol 3350, 5% isopropanol, and 0.6M Ammonium Citrate pH 5.0 at 4°C overnight. VLPs form crystals in this solution. The crystals were pelleted via centrifugation at 14,000×g for 10minutes at 4°C. The pellet was suspended in 0.3M HEPPS pH 9.0, 0.6M NaCl, and 5mM β-mercaptoethanol (β-Me) and processed through a HiPrep 16/60 Sephacryl S-500 HR (GE Healthcare Life Sciences) equilibrated with 20mM HEPPS pH 9.0, 0.6M NaCl, and 5mM β-Me. The eluted peak was concentrated using a 100kDa MWCO ultrafiltration device (Pall Corporation) and assessed with negative stained electron microscopy. Briefly, 3µl of the sample was applied to a glow discharged carbon coated 400 mesh copper grid (TedPella, 01814-F) and stained with 2% Uranyl Formate. Images were collected using a JEOL JEM-1230 microscope operating at 120kV and equipped with a Gatan US-4000 charge coupled device.

Binding studies were performed as described above. The interaction between the sorbent and 370nM concentration of VLP were measured five times to generate error bars. Statistical analysis was performed using MATLAB 2018a to generate the p-values via the Wilcoxon rank sum test (MathWorks).

## Acknowledgement

SD performed electron microscopy experiments and helped write the manuscript, BA performed electron microscopy experiments, SF performed sample preparation, RK designed and performed experiments, analyzed data, and wrote manuscript. Funds responsible for supporting these studies were provided by NIH National Institute of General medical Sciences and National institute of Allergy and Infectious Diseases (5SC1AI114843) and by Grant Number 5G12MD007603-30 from the National Institute on Minority Health and Health Disparities. Data collection for the PCV2 heparin liganded sample was performed at the Simons Electron Microscopy Center and National Resource for Automated Molecular Microscopy located at the New York Structural Biology Center, supported by grants from the Simons Foundation (349247), NYSTAR, and the NIH National Institute of General Medical Sciences (GM103310). Data collection for the unliganded PCV2 sample was performed at the Imaging Facility of City University of New York (**CUNY**) Advanced Science Research Center (**ASRC**). We would like to acknowledge the scientific and technical assistance from the NYSBC and CUNY ASRC.

The image reconstructions and atomic coordinates have been deposited into the EMDB and PDB respectively. Accession codes (EMDB/PDB) are: Unliganded image reconstruction (8939/6DZU); symmetrized liganded image reconstruction (8969/6E2R); Class 02 (8970/6E2X); Class 04 (8971/6E2Z); Class 06 (8972/6E30); Class 07 (8973/6E32); Class 09 (8974/6E34); Class 10 (8975/6E39).

## Supplemental Information

**Figure S1.**
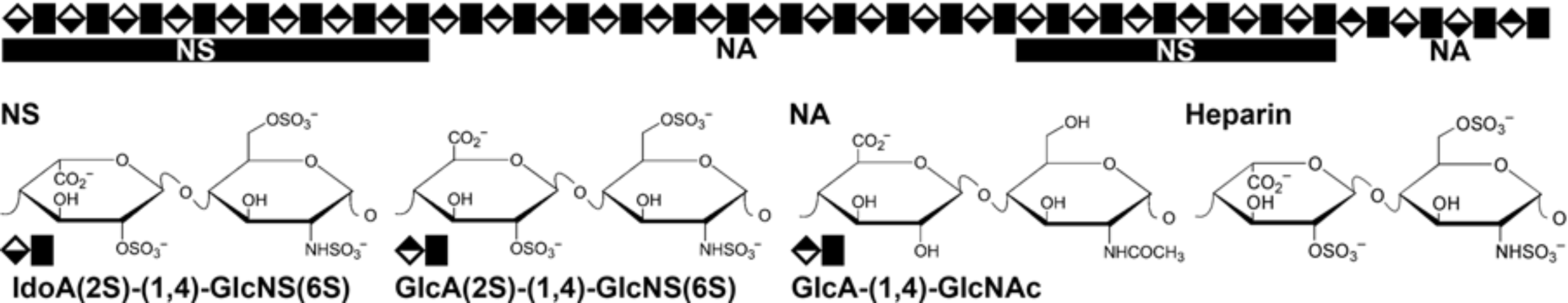
Heparan sulfate and heparin. Top) Cartoon representation of the NS and NA domains of a heparan sulfate glycosaminoglycan chain. The NS domain is composed of 3-8 repeating disaccharides, and the NA domain is composed of 2-12 repeating disaccharides. Bottom) Chemical structure of the heparan sulfate and heparin repeating disaccharides. The repeating disaccharide of the heparan sulfate NS domain is similar to the repeating disaccharide of heparin.

**Figure S2.**
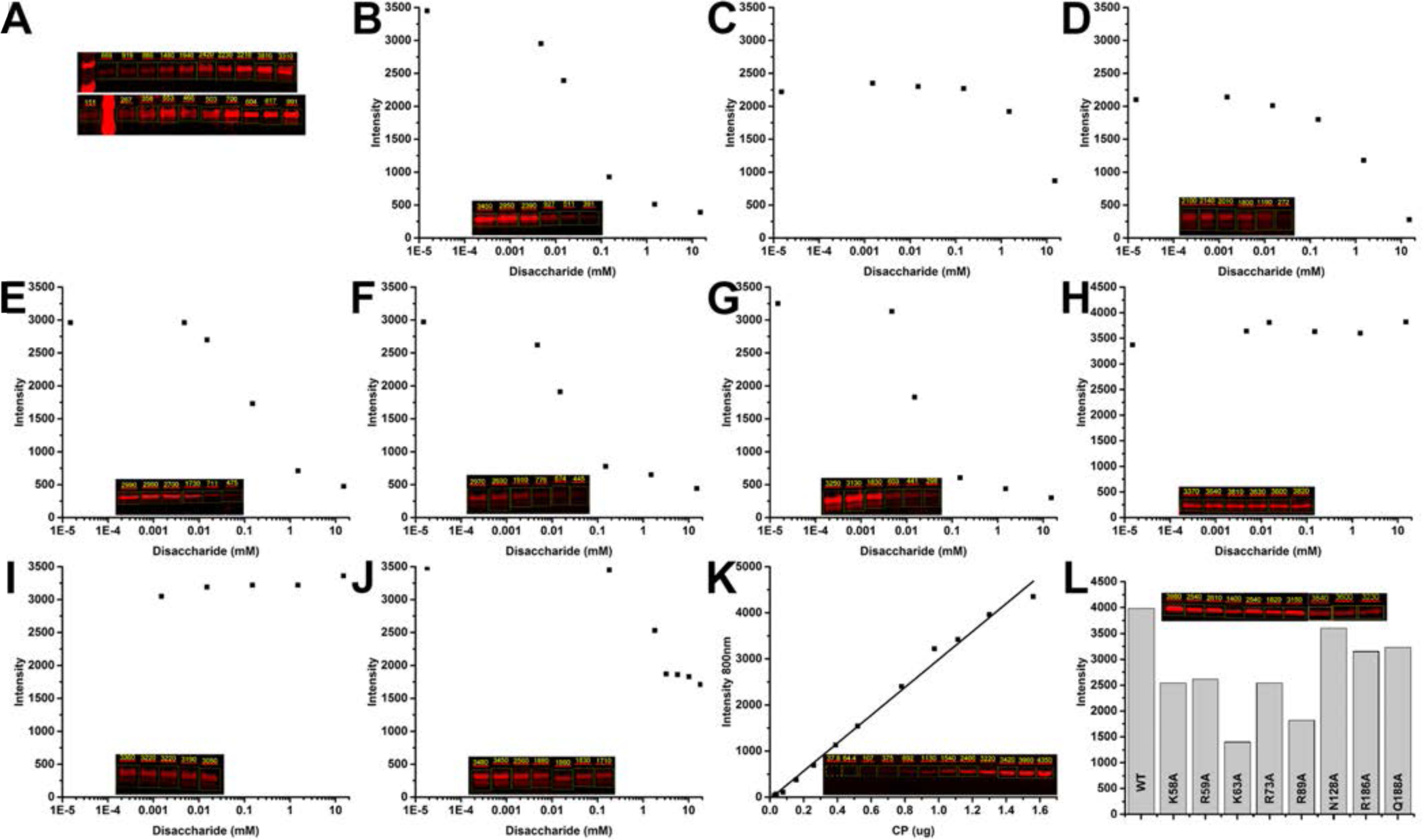
SDS-PAGE densitometry of PCV2 VLP heparin binding interaction. Images represent the interaction between PCV2 VLP and HyperD™ chromatography sorbent. The contrast of each digitized SDS-PAGE was scaled to enhance the signal for visualization. This modification does not affect the intensity reported by the Image Studio™ software used for the analysis. The numbers above each band represent the intensity corrected by background subtraction within each box. A) Binding kinetics of 370nM and 97nM VLP demonstrating that the signal from 370nM produces a robust readout (Figure 2A). B) Elution of VLP from the sorbent with increasing heparin concentration (Figure 2B). GAG competition assay representatives of: C) 6-hexose, D) 10-hexose, E) 20-hexose, F) 36-hexose heparin, G) dextran sulfate 8kDa, H) dextran 6kDa, I) hyaluronic acid, J) chondroitin sulfate B. K) Linear response curve of the LI-CORR Odyssey CLX to increasing amount of PCV2 capsid protein, indicating that intensity is directly correlated to protein content. L) Binding efficiency of *in vitro* assembled WT, K58A, R59A, K63A, R73A, R89A, N128A, R186A, and Q188A capsids (Figure 5). Inset is: WT K58A, R59A, K63A, R73A, R89A, WT, N128A, R186A, and Q188A capsids. Figures made with Image Studio™ (LI-CORR Biosciences) and Origin 2016 (OriginLab).

**Figure S3.**
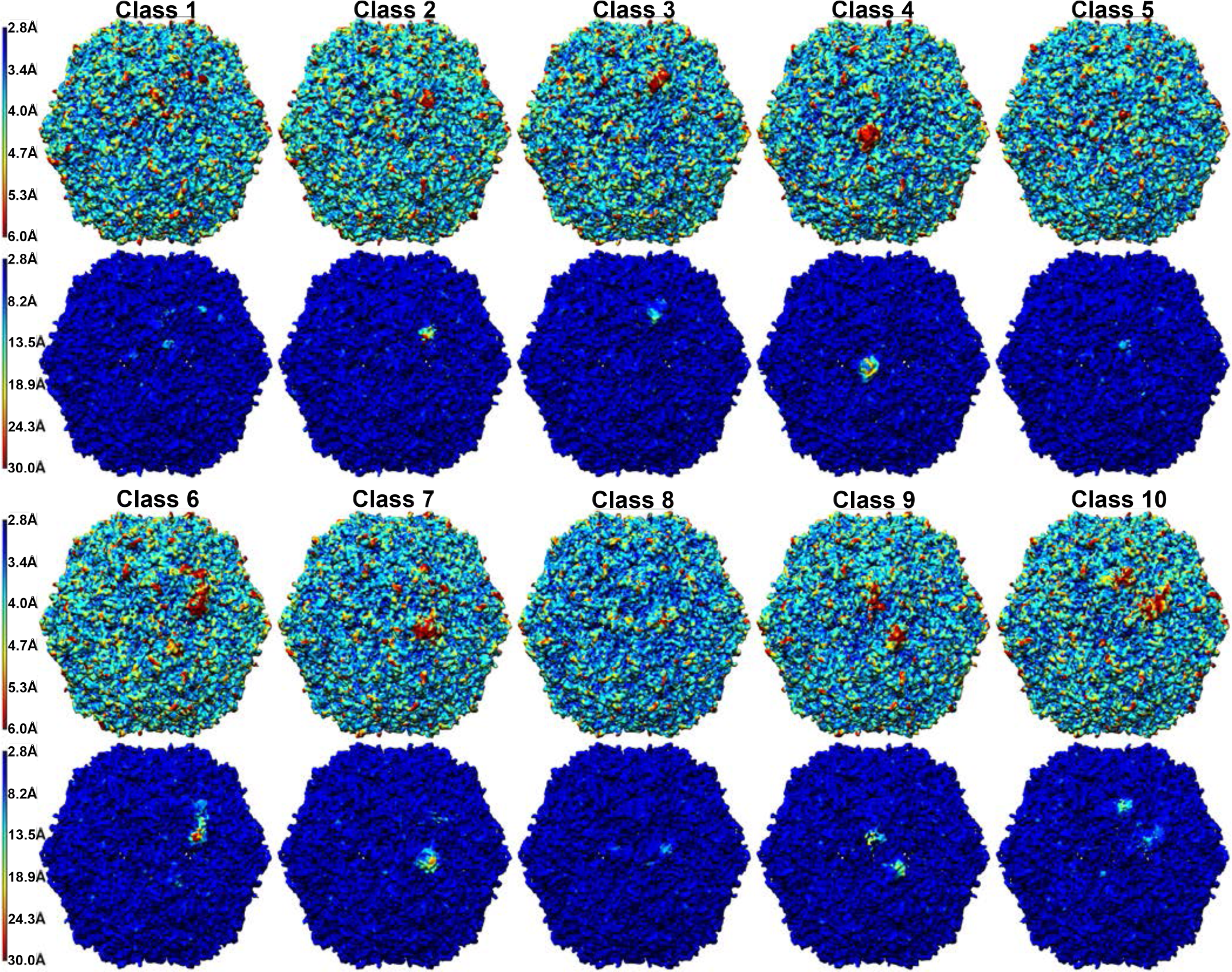
Local resolution. The local resolution for each of the 10 classes was determined using the program MonoRes. Two plots are shown for each class due to the large range in resolution. The color bar on the left of each row depicts the resolution for the indicated color. A) The top plot was calculated using a resolution range of 2.8#x212B; to 6#x212B; and the bottom plot was calculated using a resolution range of 2.8#x212B; to 30#x212B; for classes 1 to 5. B) Similar to A but for classes 6 to 10 (48). Image generated using UCSF Chimera (64).

**Figure S4.**
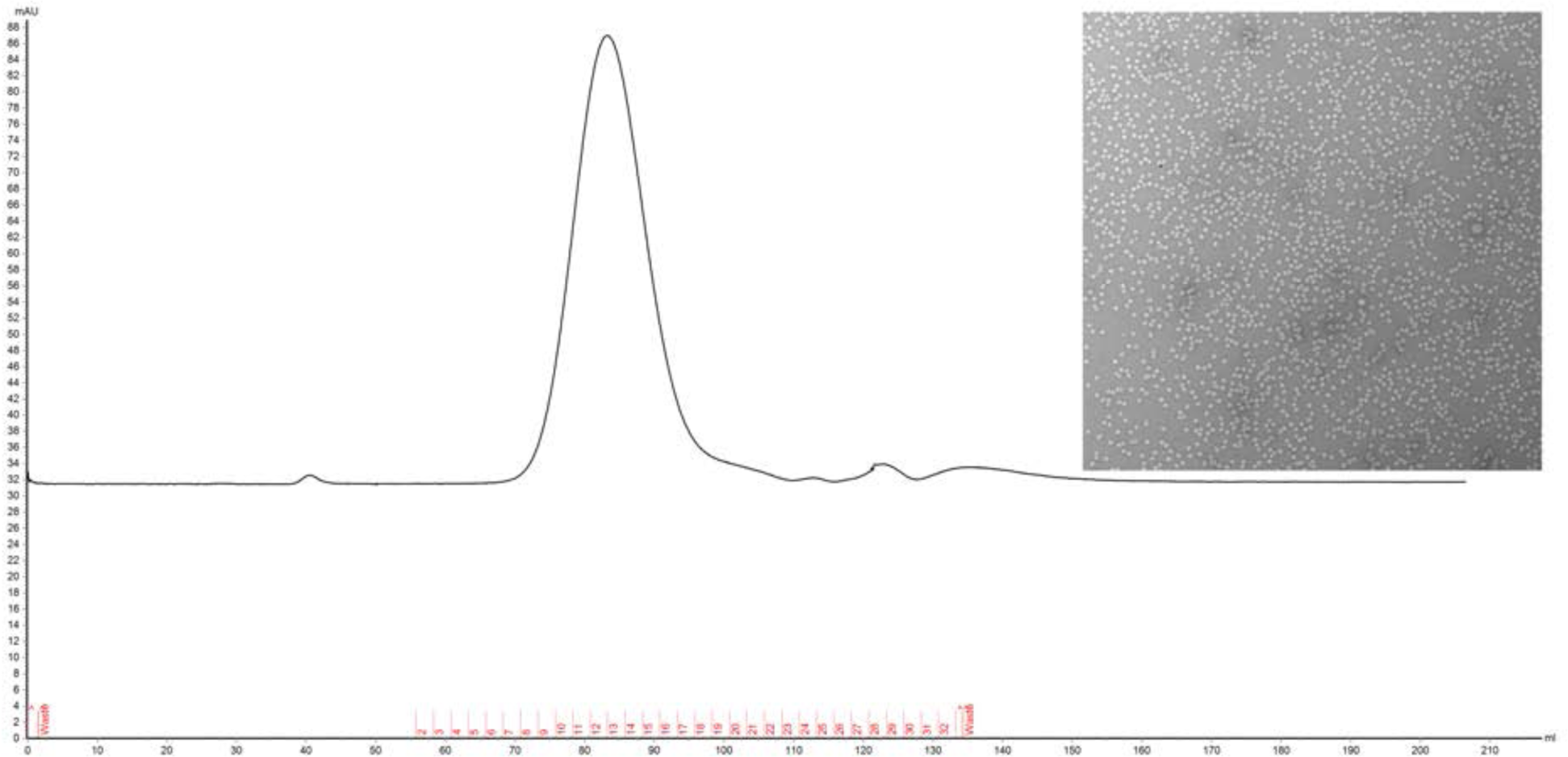
Assembly of mutant VLP. Size exclusion chromatography profile of an *in vitro* assembled PCV2 VLP. Image made with Unicorn 6.1 (GE Health Care Lifesciences) and a corresponding electron microscope micrograph of the negative stained wild-type VLP.

**Figure S5.**
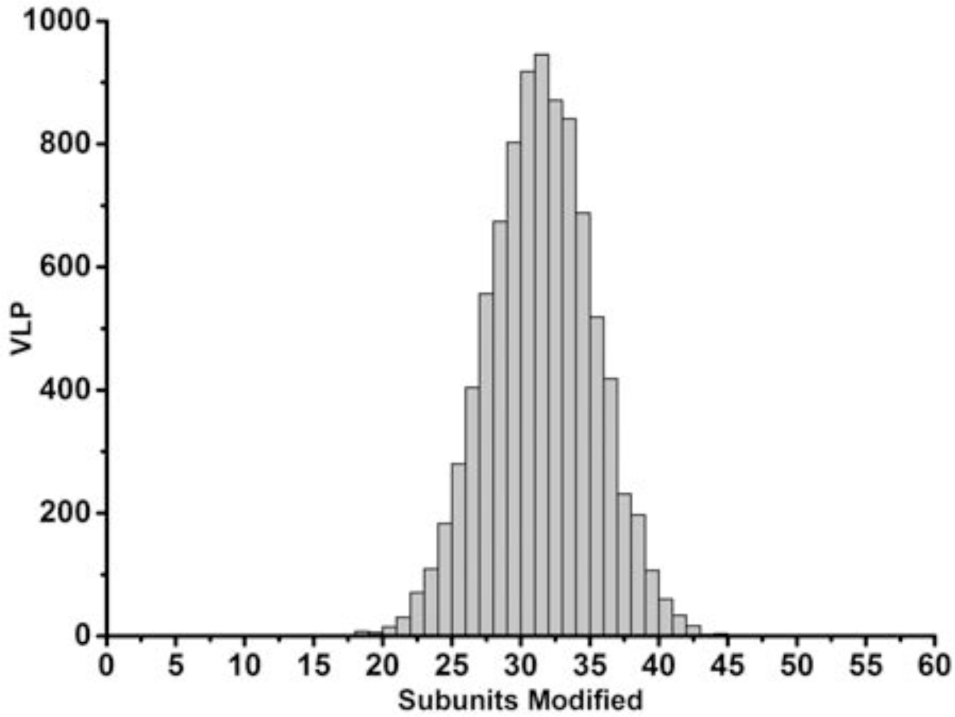
Occupied heparin binding sites on PCV2 VLPs. The distribution follows a Gaussian profile with a minimum of 17, a mode of 31, and a maximum of 44 binding sites occupied on individual VLPs. Image made with Origin 2016 (OriginLab).

